# Community successional patterns and inter-kingdom interactions during granular biofilm development

**DOI:** 10.1101/2024.01.16.575656

**Authors:** Miguel de Celis, Oskar Modin, Lucía Arregui, Frank Persson, Antonio Santos, Ignacio Belda, Britt-Marie Wilén, Raquel Liébana

**Author notes:** Corresponding authors: Raquel Liébana; Postal address: AZTI, Marine Research Division, Txatxarramendi ugartea z/g, 48395 Sukarrieta, Spain; Email address; Miguel de Celis; Postal address: Instituto de Ciencias Agrarias; C. de Serrano, 115b, 28006 Madrid, Spain.

## Abstract

Aerobic granular sludge is a compact and efficient biofilm process used for wastewater treatment which has received much attention and is currently being implemented worldwide. The microbial associations and their ecological implications occurring during granule development, especially those involving inter-kingdom interactions, are poorly understood. In this work, we monitored the prokaryote and eukaryote community composition and structure during the granulation of activated sludge for 343 days in a sequencing batch reactor (SBR), and investigated the influence of abiotic and biotic factors on the granule development. Sludge granulation was accomplished with low-wash out dynamics at long settling times, allowing for the microbial communities to adapt to the SBR environmental conditions. The sludge granulation and associated changes in microbial community structure could be divided into three stages: floccular, intermediate, and granular. The eukaryotic and prokaryotic communities showed parallel successional dynamics, with three main sub-communities identified for each kingdom, dominating in each stage of sludge granulation. Although inter-kingdom interactions were shown to affect community succession during the whole experiment, during granule development random factors like the availability of settlement sites or drift acquired increasing importance. The prokaryotic community was more affected by deterministic factors, including reactor conditions, while the eukaryotic community was to a larger extent shaped by biotic interactions (including inter-kingdom interactions) and stochasticity.

## Introduction

Aerobic granular sludge is a biofilm-based process for wastewater treatment that has received much attention in recent years. This technology displays several advantages compared to the activated sludge process, achieving an advanced nutrient removal in plants requiring about 50% less space and 20% less energy demand (Bengtsson et al., 2019; Ekholm et al., 2023). Aerobic granules are generally developed from activated sludge in sequencing batch reactors (SBRs), where aggregates with high microbial density and diversity are obtained. Substrate and oxygen gradients within the biofilm matrix allow the coexistence of ammonia-oxidizing bacteria (AOB), nitrite-oxidizing bacteria (NOB), denitrifying bacteria and phosphorous accumulating organisms (PAO), thus synchronizing nitrification, denitrification and biological phosphorus removal while degrading the organic carbon (Adav et al., 2008; Aqeel et al., 2019; Delmont et al., 2018; Wilen et al., 2018). However, there is still a lack of comprehensive studies addressing the ecological processes driving the assembly and functioning of microbial communities during granulation. There is a need to fully understand the abiotic and biotic factors influencing this process to harness the environmental and industrial potential of the technology.

Granulation is a response to specific selection pressures applied in the reactors, however, the underlying mechanisms are still poorly understood. Granules are generally obtained by 1) applying high hydrodynamic shear forces; 2) feast-famine alternation; and 3) washing-out of the non-granulated biomass (Adav et al., 2008; Aqeel et al., 2019; Wilen et al., 2018). In such reactor conditions, upon switching from planktonic to aggregated mode of growth, microbial populations ensure their persistence in flowing environments that develop under shear forces (Boltz et al., 2017). Additionally, feast-famine alternation and anaerobic feeding strategies applied in SBRs, increases bacterial cell hydrophobicity and accelerates microbial aggregation (De Kreuk and Van Loosdrecht, 2004; Liu and Tay, 2008; Wilen et al., 2018).

Eukaryotic members of the community play important roles in wastewater treatment contributing to sludge sedimentation and predation upon planktonic bacteria (Arregui et al., 2007, 2008; Burian et al., 2022a; Madoni, 2011), yet few studies have been conducted on their role in the granular sludge process. Filamentous fungi and stalked protists, have been proposed to participate in granule development by acting as a backbone for granules, thus increasing the surface to which bacteria can attach (Beun et al., 1999; Weber et al., 2007). Protistan grazing can promote aggregation of wastewater bacteria (Liébana et al., 2016), since phenotypes can switch towards biofilm development as a survival strategy (Böhme et al., 2009; Matz and Kjelleberg, 2005). But protistan grazing can also cause a reduction of bacterial biofilm thickness (Huws et al., 2005) and even extend to deep biofilm layers (Suarez et al., 2015).

The physical factors involved in granule cultivation in SBRs have been extensively studied (Liu et al., 2005; Toh et al., 2003; Verawaty et al., 2012; Wilen et al., 2018; Zhou et al., 2014) and, although to lesser extent, so has the microbial dynamics (Barr et al., 2010b; Liébana et al., 2019; Wang et al., 2009; Wittebolle et al., 2009). However, the microbial associations and their ecological implications during granular biofilm development are less understood, especially those involving inter-kingdom interactions. Here, we monitored the prokaryotic and eukaryotic community structure and dynamics during the granulation of sludge for 343 days in an SBR, aiming to elucidate the influence of abiotic and biotic factors in granular biofilm development, including their inter-kingdom interactions. For this, we studied the reactor performance and the succession of prokaryotic and eukaryotic community fractions by means of diversity and network analysis, together with null models.

## Material and methods

### Reactor set-up and operational conditions

The SBR was inoculated with activated sludge from the Hammargården wastewater treatment plant designed for biological nitrogen and phosphorus removal (Kungsbacka, Sweden) and operated at a settling time of 30 min for 343 days. The SBR, previously described in detail (Liébana et al., 2019), had a working volume of 3 L. Synthetic wastewater was used and consisted of 994.2 mg L^-1^ NaCH_3_COO, 443.8 mg L^-1^ NH_4_Cl, 139.5 mg L^-1^ K_2_HPO_4_, 56.5 mg L^-1^ KH_2_PO_4_, 12.5 mg L^-1^ MgSO_4_·7H_2_O, 15.0 mg L^-1^ CaCl_2_, 10.0 mg L^-1^ FeSO_4_·7H_2_O, and 1 mL L^-1^ micronutrient solution (Liébana et al., 2019). The feed had an organic loading rate of 2 kg COD m^-3^d^-1^, N-load of 0.3 kg NH_4_-N m^-3^d^-1^ and P-load of 0.1 kg PO_4_-P m^-3^d^-1^ resulting in a COD:N:P ratio of 20:3:1. The reactor was operated at room temperature (20-22°C) with a volumetric exchange ratio of 43%, in a 4-hour cycle of 5 min filling, 55 min anaerobic/anoxic phase, 143 min aerobic phase, 30 min settling, 5 min withdrawal and 2 min idle phase.

### Analytical methods

Concentrations in the effluent of total organic carbon (TOC) and total nitrogen (TN) were measured with a TOC-TN analyser (TOC-V, Shimadzu, Japan), and acetate, ammonium, nitrite, nitrate and phosphorus were measured using a Dionex ICS-900 ion chromatograph. The samples were immediately centrifuged, filtered (0.45 µm) and stored at -20 °C until analysis. Total suspended solids and volatile suspended solids in the reactor and in the effluent were measured according to standard methods (APHA, 1995). Microscopy was performed using an Olympus BX60 light microscope (Olympus Sverige AB, Solna, Sweden) with particle size assessment by measuring the diameter of 10 random granules using the ImageJ software (Schneider et al., 2012). A cycle study was performed on day 99. During the cycle study. a flexible plastic tube (ø 1 cm) attached to a syringe was used to sample the reactor at different heights during the aerobic phase and in the upper third of the sludge bed during the anoxic phase, to obtain representative samples.

### DNA extraction, amplification and sequencing

A total of 52 samples were collected for DNA analysis, used for both prokaryote and eukaryote amplicon sequencing analysis. DNA was extracted using the DNeasy PowerSoil Kit (Qiagen) following manufacturer’s instructions. The rDNA libraries were constructed as described in Liébana et al. (2019). Shortly, for prokaryotes, the V4 region of the 16S rRNA gene was amplified using the forward primer 515’F (5’-GTGBCAGCMGCCGCGGTAA-3’) and the reverse primer 806R (5’-GGACTACHVGGGTWTCTAAT-3’), indexed according to Kozich et al. (2013). For eukaryotes, the V9 region of the 18S rRNA gene was amplified using the 1391f (5’-GTACACACCGCCCGTC-3’) forward primer and the EukBr (5’-TGATCCTTCTGCAGGTTCACCTAC-3’) reverse primer (Amaral-Zettler et al., 2009), indexed according to Vences et al. (2016). The PCR products were sequenced with a MiSeq (Illumina) using the reagent kit v3 (PE 2x300) and v2 (PE 2x150) for the prokaryotic and eukaryotic libraries respectively.

### Sequence processing

The sequence reads were processed using the *DADA2* R version 1.22 package (Callahan et al., 2016) and *USEARCH* version 11 (Edgar, 2010), as previously described (Modin et al., 2020). The obtained count tables were used to generate consensus tables consisting of ASVs detected using both pipelines with the function *subset.consensus* implemented in *qdiv* (https://github.com/omvatten/qdiv). The taxonomic assignment was performed using the SINTAX algorithm (Edgar, 2016) based on the MiDAS database v.4.8.1 (McIlroy et al., 2015) for 16S reads and PR2 database (Guillou et al., 2013) for 18S reads. The datasets were rarefied, subsampling each sample to 43329 and 31420 reads for the prokaryotic and eukaryotic count tables, respectively. A maximum likelihood tree was generated using sequences aligned with the *msa* R package (Bodenhofer et al., 2015) and the *phangorn* R package (Schliep, 2011) was used to fit a GTR + G + I maximum likelihood tree. Taxonomic α-diversity was calculated using Hill numbers (Jost, 2006) using the *hillR* R package (Li, 2018). The effect on α-diversity of biological (α-diversity of the other group) and environmental parameters was evaluated using linear models. The Hill numbers framework was also used to calculate β-diversity (Modin et al., 2020), dissimilarity indices (^q^βdis) constrained between 0 and 1 using *qdiv*. Community succession and its relationship with environmental parameters were evaluated performing distance-based Redundancy Analysis (dbRDA) and variance partitioning analysis using Bray-Curtis dissimilarity matrices with the vegan R package (Oksanen et al., 2019).

### Network analysis

Network analysis was conducted to evaluate the interaction patterns of the bacterial and eukaryotic communities within the bioreactor. We first removed the ASVs present in less than 10% of samples and with an abundance lower than 0.1% (resulting in 411 and 125 ASVs remaining in the prokaryotic and eukaryotic datasets respectively). Then, we calculated every potential co-occurrence between the ASVs applying two correlation models, Spearman’s rank correlation and Sparse Correlations for Compositional data (SparCC), implemented in the *SpiecEasi* R package (Kurtz et al., 2015). Co-occurrence were considered when the Spearman’s correlation coefficient (ρ) and SparCC R-corr absolute values were higher than 0.6, and their false discovery rate (FDR) corrected p-values lower than 0.05. The resulting networks consisted of 325/67 nodes and 6010/308 edges for the prokaryotic and eukaryotic communities, respectively. Co-occurrence patterns of the core communities and the potential interkingdom associations were assessed on filtered networks, keeping the nodes present in more than 75% of samples. Network visualization and nodes’ module membership calculation were performed with the *igraph* R package (Csardi and Nepusz, 2006). We also calculated the proportion of ASVs (module completeness) and the abundance of each assigned module in the networks. In addition, we applied the method developed by Ortiz-Álvarez et al. (2021) to calculate the individual co-occurrence networks of each time-step sample, to calculate their individual properties and assess the structure of the microbial communities over time.

### Microbial community phylogenetic dispersion against a null expectation

We assessed the influence of stochastic and deterministic processes in the community succession by means of null model analysis on the within (α) and between (β) sample phylogenetic diversity, coupled with taxonomic turnover (Supporting Material). Briefly, we first tested the phylogenetic signal of the communities, that is, if closely related ASVs have similar environmental preferences, allowing us to relate phylogenetic dispersion with ecological processes driving community succession (Stegen et al., 2012). We used α-diversity-based null models to calculate the phylogenetic dispersion of the communities, accounting for the whole phylogeny or just the tip of the tree (Webb et al., 2002). Thus, we calculated the net relatedness index (NRI) and nearest taxon index (NTI) using the *ses.mpd* and *ses.mntd* functions of the *picante* R package (Kembel et al., 2010). The closer they get to zero, the closer the phylogenetic structure of the community is to the null expectation, reflecting the higher influence of stochasticity (Webb et al., 2002). Phylogenetic β-diversity was used to evaluate the ecological process dominating community succession over time, comparing the phylogenetic dispersion between two consecutive samples. For this, the β-Nearest Taxon Index (βNTI) was calculated with the *qdiv* package (Modin et al., 2020). Contrary to NRI/NTI, positive values of βNTI reflect phylogenetic overdispersion, whereas negative values indicate phylogenetic clustering. In summary, |βNTI| > 2 is indicative of a deterministic process that has caused two microbial communities to be either phylogenetically clustered (βNTI < -2) or dispersed (βNTI > 2). Taxonomic turnover was assessed using Raup-Crick based measures, calculated using the *qdiv* package, which quantify the deviation of the observed turnover from that expected if the community was randomly assembled. |RC_bray_| values > 0.95 are considered to reveal that the observed community composition is different from the null expectation, whereas |RC_bray_| values < 0.95 are consistent with the effect of drift (Stegen et al., 2013).

## Results

### Different stages identified during granulation

High wash-out rate with reactor operation at short settling times (typically 1-5 minutes for aerobic granular sludge systems operated at lab-scale) has been claimed as a prerequisite for granulation to occur (Morgenroth et al., 1997). In this study, granulation was observed at long reactor settling time of 30 minutes (Supplementary Figure S1, Supplementary Figure S2). Granules started to emerge at day 16 and the mean particle size increased, especially after day 115, once the granules were completely developed (Supplementary Figure S2). According to the granulation process, we identified three stages: floccular stage (days 0-15), intermediate stage (days 16-115) and granular stage (days 116-343). The sludge concentration, with a median volatile fraction of 77% (SD = 12), and the sludge retention time increased as particle size did, especially in the granular stage (Figure 1A, B), while the effluent suspended solids concentration was generally below 50 mg L^-1^ (Figure 1C). Carbon removal was stable during the experiment (Figure 1D), with TOC removal generally above 97%. Complete ammonium removal was achieved during most of the experiment (Figure 1E), showing a median removal of 97% (SD=15). Nitrification occurred in the reactor as nitrite was mostly not present in the effluent and nitrate was formed (Figure 1F, G), especially once granules emerged. The median total nitrogen removal was 48% (SD=23), being variable along the experiment (Figure 1D). Denitrification took place in the reactor as the depletion of nitrate within the SBR cycle was observed (Supplementary Figure S1). Biological phosphorus removal occurred in the reactor (Supplementary Figure S1) but the removal was variable and had an increasing trend with time (Figure 1H).

**Figure 1.**
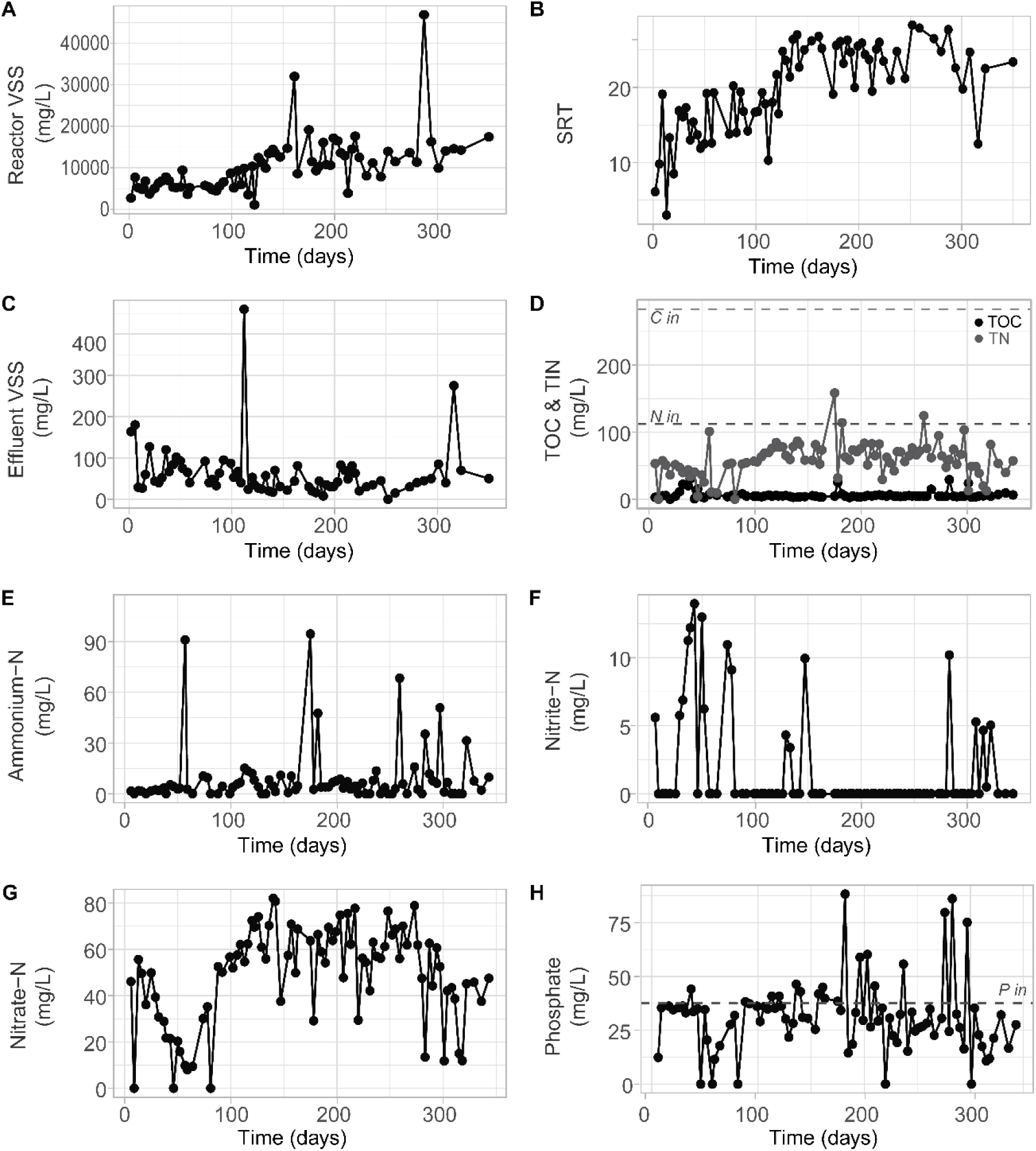
Sludge and performance parameters during the reactor run. A, reactor volatile suspended solids (VSS) concentrations; B, sludge retention time (SRT); C, effluent volatile suspended solids concentrations; D, in black, effluent total organic carbon (TOC) and in grey, total inorganic nitrogen (TIN) expressed as the addition of ammonium, nitrite and nitrate; E, effluent ammonium concentration; F, effluent nitrite concentration; G, effluent nitrate concentration; H, effluent phosphate concentration. Horizontal dashed lines indicate the influent concentration: total organic carbon 283 mg L-1 (C in), total nitrogen 112 mg L-1 (N in) and phosphorous 37.6 mg L-1 (P in).

### Pronounced shift of dominant taxa during the early stages of granulation

Overall, we observed marked compositional changes during the floccular and intermediate stages while the granular stage was characterized by being more compositionally stable in both eukaryotic and prokaryotic community fractions. The microbial community composition displayed drastic shifts after the reactor start-up with replacement of initially dominant taxa (Figure 2). The initial prokaryotic community was complex. The genera *Acinetobacter*, *Thauera* and *Dechloromonas* had high relative abundance and the latter increased during the floccular stage. During the intermediate stage, *Candidatus Accumulibacter* and *Zoogloea* increased in abundance, and later dominated the granular stage, together with *Defluviicoccus*, *Ferribacterium* and *Rubrivivax*. These changes were also evident for the eukaryotic community. During the floccular stage, the microeukaryotic community was represented by members of the SAR supergroup, dominated by ASVs affiliated to *Pseudofungi* (*Oomycota*), *Sagenista* (*Fibrophrys*) and *Oligohymenophorea* (*Sessilida*) groups. These, were rapidly replaced by the *Roghostoma* genus (*Rhizaria* group) which decreased when granules started to dominate, with an accompanying transitional increase in abundance of the rotifer *Platyias* sp. Finally, during the granular stage, ASVs affiliated to *Rhogostoma* sp. kept their presence, dominating the eukaryotic community, accompanied by members of *Cryptomonadales* (*Hacrobia*), *Opisthokonta* (*Rotifera*) and *Alveolata* (*Sessilida*) (Supplementary Figure S6).

**Figure 2.**
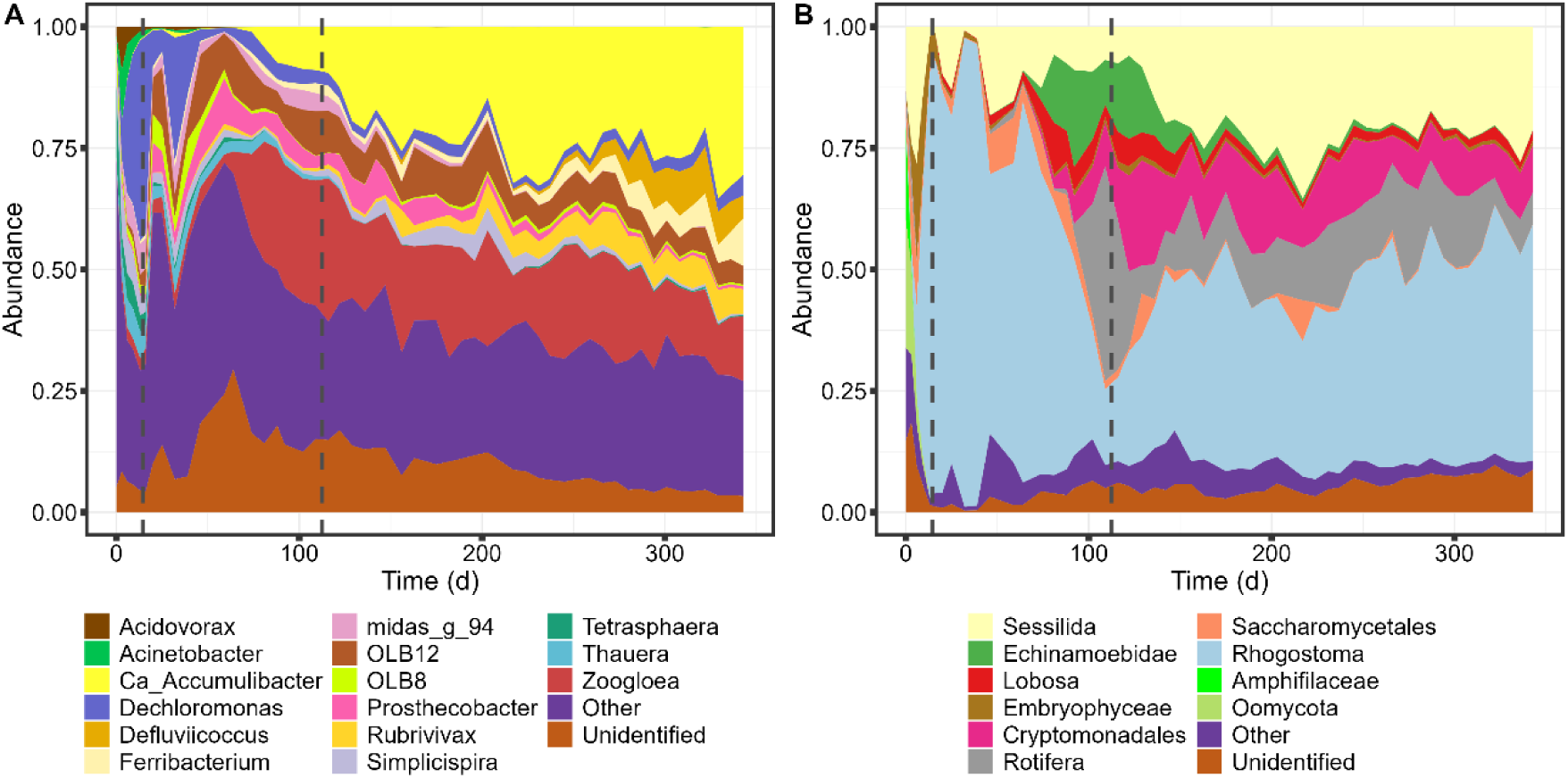
Temporal dynamics of microbial taxonomic composition. Taxonomic distribution of A, prokaryotic community at the genus-level and B, eukaryotic community at the family-level abundances. “Other”, includes minor prokaryotic genera and eukaryotic families; and “Unidentified”, taxonomically unassigned taxa.

We also evaluated eukaryotic communities through microscopical observations. Due to their dimensions and morphological characteristics, *Sessilida*, sessile peritrich ciliates, were the microeukaryotes most easily observed (Supplementary Figure S2). They exhibit a sessile stage fixed by a stalk to a substrate as well as a dispersal stage with free-swimming forms that seek new substrates during their life cycles (Gilbert and Schröder, 2003). Species with a morphology compatible with the *Epistylis* genus predominated throughout the study. In the floccular stage, cells with *Vortecellides*-like morphology, having a stalk with spasmoneme, were also observed. *Epistylis* is a colonial genus with a thicker, rigid, not contractile stalks, more adapted to rapid water currents (Taylor, 1983), the attachment of other sessile filter feeders seem to be inhibited during the process of granulation. Colonies with a varying number of zooids and stalks lengths (indicating a probable coexistence of different species within the genus) were distributed throughout the entire surface of the granules although some areas seem to be more favorable to the attachment than others. *Rhogostoma* are small testate amoebas that dominated mainly during the intermediate stage, and were also dominant during the granular stage (Figure 2B), but which direct observation is not as common as peritrichs. These protists are raptorial feeders (Parry, 2004) with pseudopods which allow searching for bacterial prey that are loosely associated or permanently attached to surfaces.

### Parallel prokaryotic and eukaryotic community succession

Both bacterial and eukaryotic communities suffered a similar drastic decrease in α-diversity during the first days of operation, with and without accounting for relative abundance (Figure 3A-C). During this period, the taxonomic α-diversity of the bacterial community dropped by around 50% for ^0^TD and by over 60% for ^1^TD and ^2^TD, while the eukaryotic community showed even a higher loss, over 65% for ^0^TD and over 85% for ^1^TD and ^2^TD. At the beginning of the intermediate stage, the prokaryotic community increased its diversity, followed by a decreasing trend by the end of this stage and stabilization by the end of the experiment. The eukaryotic community increased in abundance by the end of the intermediate stage, followed by a decreasing trend throughout the granular stage. The loss of diversity over the experiment was more pronounced for the prokaryotic community than the eukaryotic. These trends were more pronounced when accounting for the dominant ASVs (i.e., when q = 2).

**Figure 3.**
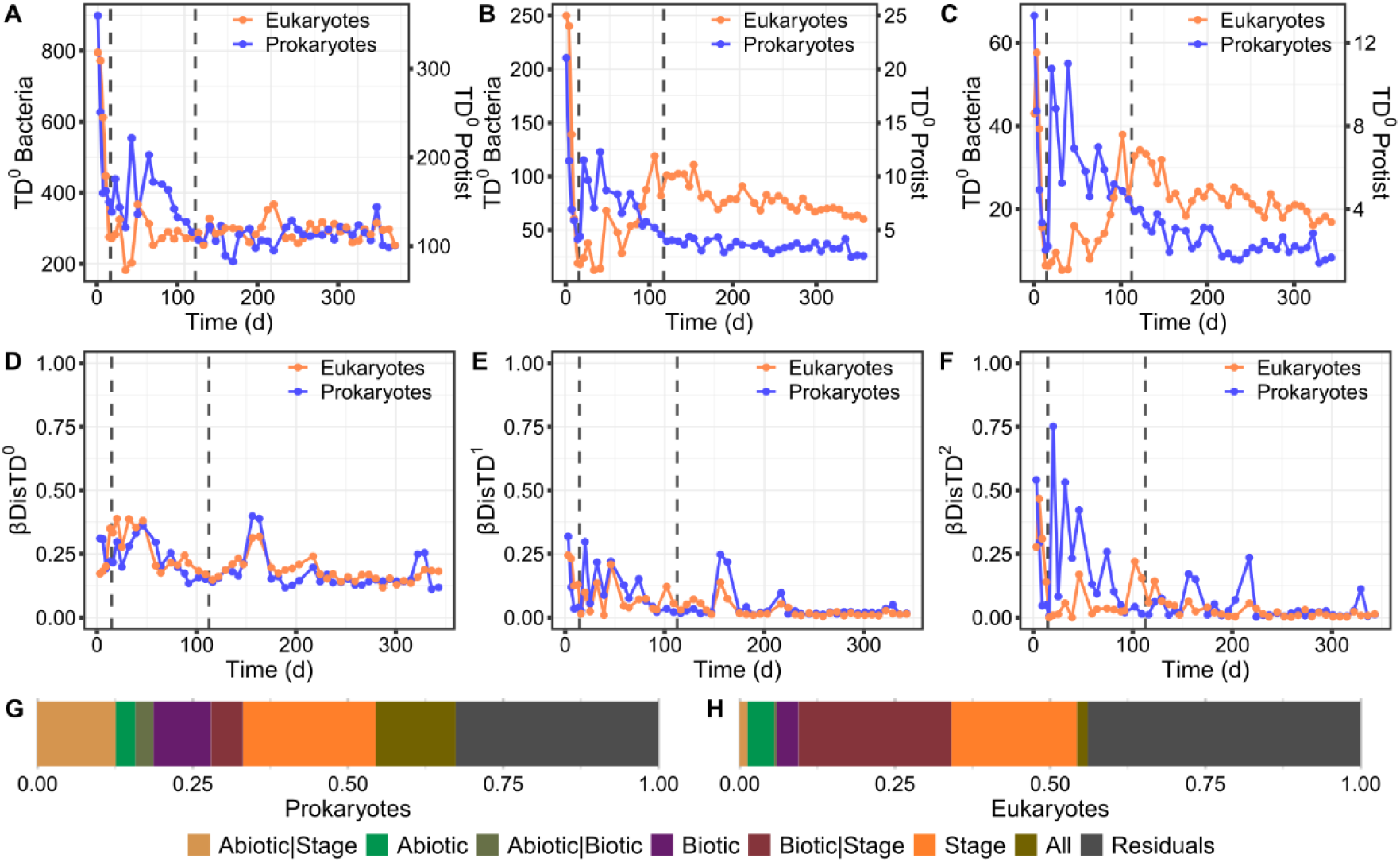
Microbial diversity and succession patterns. A-C, α-diversity (qTD) based on Hill numbers, as a function of order q. D-F, β-diversity (βDisTDq) between two successive sample points of the prokaryotic and eukaryotic communities. A, D, q of 0, relative abundance is not considered for the calculation. B, E, q of 1, relative abundance is considered. C, F, q of 2 more weight given to more abundant ASVs. Vertical dashed lines represent the three stages of biomass granulation in the bioreactor: Stage 1 – Flocs; Stage 2 – Intermediate; Stage 3 – Granules. Variance Partitioning Analysis describing the percentage of G, prokaryotic and H, eukaryotic community variation explained by sample data categories (Abiotic – sludge and performance parameters, Biotic – α-diversity of the other group, Stage – granulation stage of the reactor at each sample point), and their combined effect (e.g., Abiotic | Biotic). Variation not explained (residuals) and shared variation between the three categories are also shown.

Prokaryotic and eukaryotic community succession patterns were parallel over the whole experiment (Figure 3D, E), especially when accounting for rarer ASVs (i.e., q = 0 and q = 1; Supplementary Table S1). We observed fewer changes between successive communities at the end of the experiment, as reflected by taxonomic composition and α-diversity. In addition, the dominant prokaryotic community (q = 2) underwent more changes during the intermediate stage (Figure 3F), in line with the taxonomic shifts observed (Figure 2A). Both eukaryotic and prokaryotic communities presented parallel community succession, as revealed by constrained ordination and β-diversity correlation (Figure S3). Variance partitioning analysis revealed that biotic factors were more important for the eukaryotic community (Figure 3H), whereas the prokaryotic community was more affected by abiotic factors such as nitrate or suspended solids concentration (Figure 3G). Both succession trends were determined by the diversity of the other trophic level. The prokaryotic succession was significantly affected by the increase in diversity of dominant eukaryotes by the end of the intermediate stage. In the case of the eukaryotic succession, the higher prokaryotic diversity at the beginning of the experiment significantly affected the succession.

### Network analysis reveals changes in community structure during granule formation

Both prokaryotic and eukaryotic metawebs were divided in four modules, or sub-communities, with parallel dynamics when attending to their proportion over time. Prok-3 and Euk-3 dominated the prokaryotic and eukaryotic communities in the initial floccular phase. These modules were replaced by Prok-1 and Euk-1 in the intermediate phase, and subsequently by Prok-2 and Euk-2, which dominated in the granular phase (Figure 4AB; Supplementary Figure S4). Time-point network properties revealed differences between prokaryotic and eukaryotic dynamics of community structure (Figure 4C-D), although both communities presented less clustering during the intermediate stage. Modularity and edge density followed opposite trends in both communities. In the prokaryotic community, modularity increased during the intermediate stage and decreased during the granular stage. The eukaryotic modularity was at a minimum during the first granulation stages and increased starting the granulation stage.

**Figure 4.**
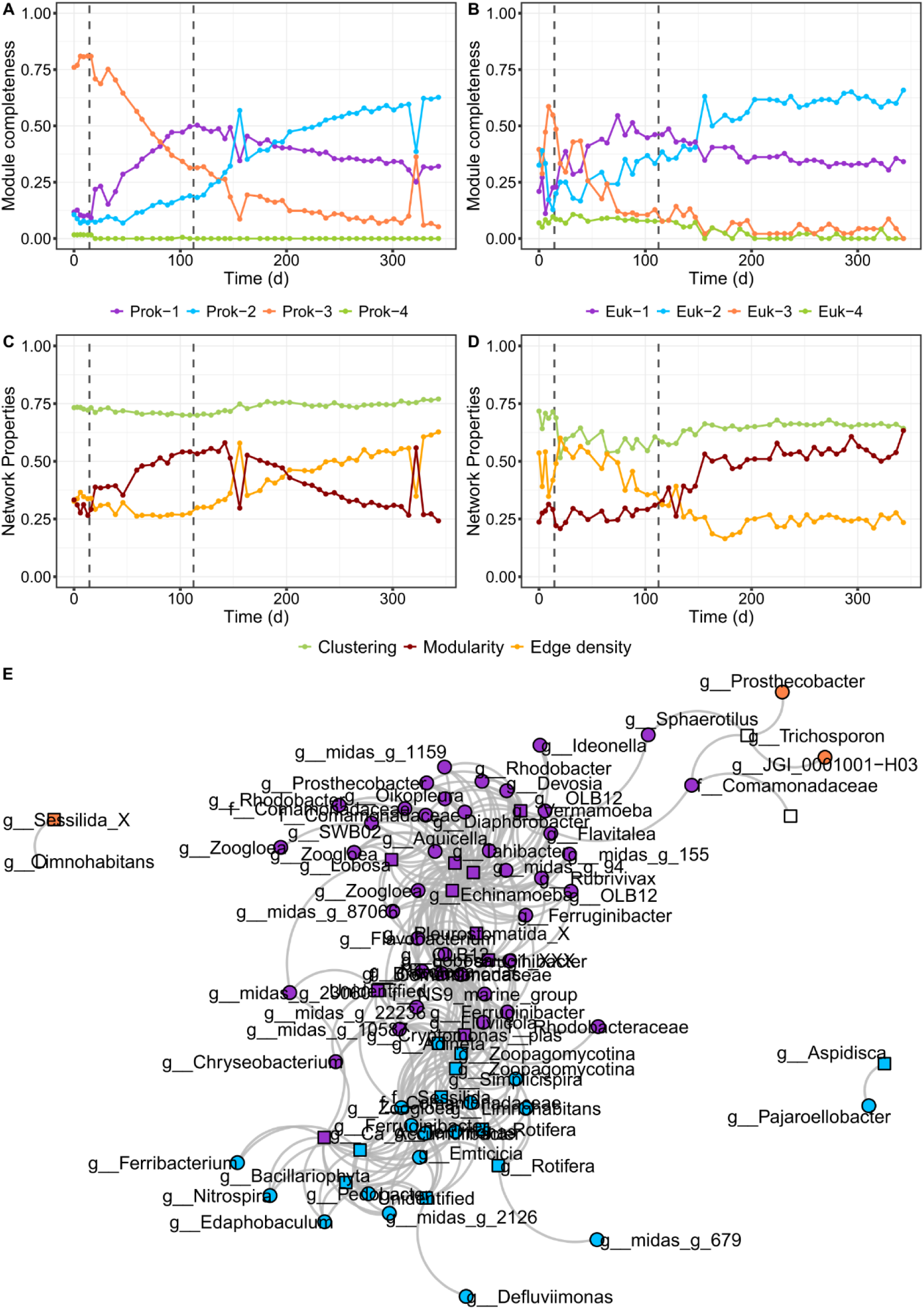
Composition and interaction structure dynamics of the microbial communities in the bioreactor. Community composition structure measured as the proportion of nodes belonging to a given module (i.e., module completeness) of the A, prokaryotic and B, eukaryotic communities. Temporal patterns of modularity, clustering coefficient, and edge density of the C, prokaryotic and D, eukaryotic communities. E, Bipartite network showing the significant positive correlations between prokaryotic and eukaryotic core ASVs. Vertical dashed lines represent the three stages of biomass granulation in the bioreactor: Stage 1 – Flocs; Stage 2 – Intermediate; Stage 3 – Granules.

We further investigated the main potential microeukaryote-prokaryote interactions in the reactor using bipartite networks. Thus, we calculated the eukaryote-prokaryote co-occurrence networks keeping only the nodes that were detectable through most of the experiment (>75% samples). Besides, we used the module membership of the previously calculated networks to assess the potential interactions between sub-communities with similar behaviour (Figure 4E). In this network, the nodes closely connected belonged to the prokaryotic and eukaryotic modules behaving similarly (i.e., nodes from Prok-1 and Euk-1, or nodes from Prok-2 and Euk-2). Throughout the granulation stage, Prok-2 and Euk-2 sub-communities dominated the microbial communities, with an abundance of 55.3% and 29.1% by the end of the experiment. However, they were simpler, with fewer nodes and connections than Prok-1 and Euk-1, which were the dominant communities during the intermediate stage (Figure 4, Supplementary Table S2). The most abundant ASVs from Prok-2 belonged to *Ca.* Accumulibacter (30.4%), *Ferribacterium* (9.9%) and *Zoogloea* (9.8%) genera, and from Euk-2 to the *Sessilida* (21.0%) family and the *Rotifera* (5.4%) (Supplementary Figure S4).

### Prokaryotic and eukaryotic community succession is governed by different ecological processes

We assessed the phylogenetic alpha dispersion of the microbial communities over time by calculating the net relatedness index (NRI) and the nearest taxon index (NTI), which examines the clustering/dispersion of phylotypes (Figure 5A, B). First, to justify the use of null models on phylogenetic α and β diversities, we verified the phylogenetic signal across relative short phylogenetic distances (Supplementary Figure S5). We observed that the NRI (prokaryote: 0.59 ± 1.10; eukaryote: -0.36 ± 1.55) values were lower than the NTI (prokaryote: 3.69 ± 0.76; eukaryote: 1.30 ± 1.08) values in both communities revealing that deterministic assemblage is more relevant at terminal levels in the phylogeny. Besides, we found that the NRI values were much higher in the first samples, and then decreased. In the prokaryotic community the higher NRI values match with the floccular stage (NRI = 2.23 ± 0.68), and then was at some extent stable around 0 (0.41 ± 0.99). The NRI values of the eukaryotic community decreased faster during the first samples, reaching values close to 0 in the intermediate stage and negative values in the granular stage. This decreasing trend is also observed in the eukaryotic NTI, reaching values close to 0 at the end of the experiment. The NTI of the prokaryotic community, however, remained positive during the whole experiment.

**Figure 5.**
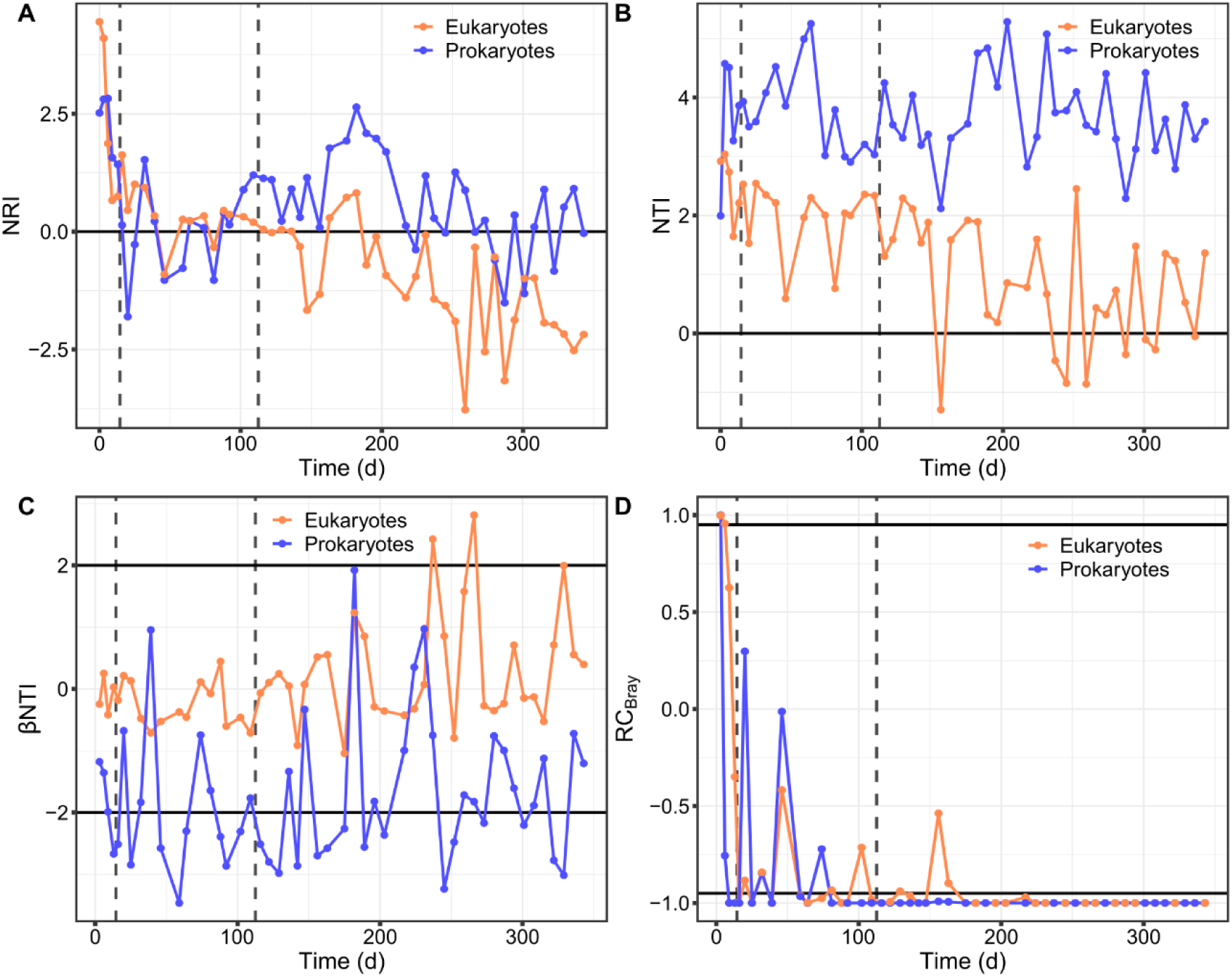
Temporal changes in phylogenetic structure and successional turnover in the bioreactor. Community phylogenetic structure assessed via A, net relatedness index (NRI) and B, nearest taxon index (NTI). Null model analysis results between two successive sample points assessed by C, phylogenetic turnover based on βNTI, and by D, taxonomic turnover based on RCBray. Horizontal lines indicate thresholds for significant deviations from the null expectation. Vertical dashed lines represent the three stages of biomass granulation in the bioreactor: Stage 1 – Flocs; Stage 2 – Intermediate; Stage 3 – Granules.

Then, we evaluated the changes in β-diversity using the RC_bray_ (based on taxonomic turnover), and βNTI (based on phylogenetic turnover) metrics (Figure 5C, D). The RC_bray_ values were higher than the null expectation (RC_bray_ > 0.95), then the values decreased in both communities, however, their dynamics differed. In both, the prokaryotic and eukaryotic communities, RC_bray_ values were within the null expectation (|RC_bray_| < 0.95) in the initial and intermediate granulation stages, and in the granular stage were overall lower than expected (RC_bray_ < -0.95). The βNTI dynamics also differed between prokaryotic and eukaryotic communities. The prokaryotic community had overall negative βNTI values, mostly lower than the null expectation (βNTI < -2). However, the eukaryotic community had βNTI values around 0, and by the end of the experiment some values were higher than the null expectation (βNTI > 2).

## Discussion

To date, there is still a lack of comprehensive studies addressing the microbial associations and their ecological implications occurring during granule development in SBRs, especially those involving inter-kingdom interactions, as relatively few studies have been conducted on the role of eukaryotes on the granular sludge process. Here, we studied the microbial community succession during sludge granulation. Sludge granulation has been described to occur in several steps (Liu and Tay, 2002) including cell-to-cell contact and micro-aggregation, maturation of the microbial aggregates by forming a matrix of extracellular polymeric substances (EPS) where cells attach and multiply, and granule size increase and microbial stratification within the granular matrix (Wilen et al., 2018). In this work, during sludge granulation we indeed identified three ecological stages, floccular, intermediate and granular, when both the microbial community and sludge parameters changed.

### The initial granulation stage is characterized by rapid community shifts

The initial stages of the granulation process are controlled by different environmental factors and properties of the biomass (Liu et al., 2009; Liu and Tay, 2002), which are closely related with the microorganisms inhabiting and dominating the bioreactors. We found the prokaryotic community dominated by bacteria (e.g., *Acinetobacter* sp., *Dechloromonas* sp. or *Thauera* sp.) which have been described as early biofilm colonizers and important EPS producers (Gao et al., 2022; Katharios-Lanwermeyer et al., 2014; Liébana et al., 2016) and commonly detected during the initial phases of granulation (Liébana et al., 2019; Weissbrodt et al., 2012; Xia et al., 2018). A diverse community of microeukaryotes was also present, mostly protozoa with different feeding modes and motility which promote microbial activity and aggregation against predation (Yang et al., 2022). The *Rhogostoma* genus increased in abundance, dominating the community by the end of the floccular stage. They have been observed as dominating eukaryotic communities in several WWTPs (Chouari et al., 2017; Hirakata et al., 2019; Matsunaga et al., 2014; Remmas et al., 2017) and found to be represented by a single species, *Rhogostoma minus,* which recently have received researchers attention not only for its wide spread distribution but for hosting well-known human pathogenic *Legionellales* (Pohl et al., 2021). The abundance of the *Sessilida* family (*Alveolata* supergroup) in turn reached a minimum at the end of the floccular stage. This could be explained by a sudden increase of rotifer populations, that could be ingesting *Sessilida* individuals (Li et al., 2013). On the other hand, it may be related to the decrease of available preys (Johansson et al., 2004), as a result of the prokaryotic species filtering occurring in the reactor due to the acclimation of the sludge community to the new environmental conditions, evidenced by the pronounced drop in microbial α-diversity (with and without accounting for relative abundance). These changes were also revealed by the community successional patterns, indicating a higher turnover in community composition during the first stages of granulation, especially when accounting for the relative abundance. The switch from complex to simple and easily biodegradable substrate (i.e., acetate) could explain the drop in prokaryotic α-diversity and the higher community dynamics as a result of strong selective processes during the first stage of granulation (Liébana et al., 2019; Szabó et al., 2017; Weissbrodt et al., 2012). Consequently, the changes in the prokaryotic community structure would also influence the diversity and abundance of heterotrophic eukaryotes (Burian et al., 2022b; Saleem et al., 2013). Indeed, module completeness revealed the division of prokaryotic and eukaryotic communities into sub-communities with, presumably, different environmental preferences (de Celis et al., 2022). During the floccular stage both were dominated by a disappearing sub-community (Prok-3, Euk-3), allegedly adapted to the initial activated sludge conditions. Prokaryotic and eukaryotic initial communities had increased NRI and NTI values revealing phylogenetic clustering, previously observed in activated sludge systems (de Celis et al., 2022), which could be indicative of the deterministic forces driving community assembly.

During sludge granulation, the suspended biofilms further develop and increase in diameter where oxygen and substrate gradients are created within the granule, providing new niches to be colonized. Indeed, when granules started to emerge during the intermediate stage, the microbial community displayed high fluctuations. During this stage, important EPS producers such as *Zoogloea* sp. (Seviour et al., 2012) and others associated with the production of a resistant matrix of structural EPS like *Ca. Accumulibacter* (Guimarães et al., 2023) substantially increased in abundance. Members of the *Rhogostoma* protistan group decreased in relative abundance in favor of other micro-eukaryotes within the *Cryptomonadales*, *Rotifera* and *Sessilida* groups. The areas with greater abundance of peritrichs (*Sessilida*) were those with irregularities and grooves as they can find protection against water turbulence and hence, less possibilities of getting detached (Sartini et al., 2018) although signals of abrasion, i.e. lack of zooids, were frequently observed. The denser peritrichous colonization during this stage (Supplementary Figure S2) may reflect the increased availability of settlement sites, and the favorable conditions for the development of these bacterivore ciliate communities (Madoni, 2005; Patterson and Simpson, 1996). Additionally, the flourish of stalked ciliates could have improved the granulation process serving as backbone for biofilm development (Weber et al., 2007).

### Granule development allows for community stabilisation

As the granulation progressed, the prokaryotic community stabilized at the beginning of the granular stage, observing a replacement of the initial community by a simpler prokaryotic community dominated by a handful of bacterial genera including Ca*. Accumulibacter*, *Defluviicoccus*, *Ferribacterium*, *Rubrivivax* and *Zoogloea*, commonly detected in reactors performing simultaneous biological removal of organics, nitrogen and phosphorous in wastewater treatment plants (Dris et al., 2015; Lemaire et al., 2008; Winkler et al., 2018). We indeed observed nitrification and phosphorous removal to improve during this phase. In agreement with the progression of the granulation, the biomass concentration in the reactors was doubled. The granular size increase changed the microenvironment within the granule matrix, thus increasing the importance of deterministic factors influencing the community assembly process during this stage, both when considering the terminal levels in the phylogeny (NTI), or the taxonomic turnover between successive samples (RC_Bray_). The phylogenetic turnover between successive prokaryotic communities (βNTI) also indicated the importance of deterministic factors driving community succession. This agrees with the observed higher influence exerted by abiotic factors, such as nitrate concentration in the prokaryotic community. The eukaryotic community succession was more affected by biotic factors, such as inter-kingdom interactions, competition, predation and mutualism (Bock et al., 2020) or random factors such as free settlement sites for reproduction, explaining the overall lower values of NRI and NTI and overall random phylogenetic turnover, compared to prokaryotic communities. In addition to trophic interactions, eukaryotic and prokaryotic communities also compete for physical niche. For example, due to their similar growth pattern filamentous bacteria compete for settling sites with peritrichous ciliates (Stoessel, 1989), which could contribute to the opposite trends in prokaryotic and eukaryotic richness during early granule formation.

Network analysis also evidenced the changes associated by granule particle size increase. A higher taxonomic turnover was revealed by emerging sub-communities (Prok-1, Euk-1; Supplementary Figure S4). The presence of differently sized aggregates promoted the generation of different niches for functional groups (Liu et al., 2020), like the dominating Prok-1 sub-community (members of the *Comamonadaceae* family and *Zoogloea* sp.). This niche differentiation process could also be revealed by the initial increase in prokaryotic modularity coupled with a decrease in clustering coefficient (Ortiz-Álvarez et al., 2021). The eukaryotic community presented a similar increasing modularity trend by the end of the intermediate stage, when peritrichs of the *Sessilida* family and numerous *Rotifera*, within the Euk-2 sub-community, emerged (Supplementary Figure S2, Supplementary Figure S4). Besides, this stage was also characterized by a stabilization of a *Cryptomonas* population (from Euk-1), which would not compete with the Euk-2 sub-community for settling sites (Salmaso and Tolotti, 2009). *Cryptomonas* is a mixotrophic genus with species that can combine photosynthetic activity with utilisation of exogenous carbon sources, here, uptake of supplemented acetate or/and engulfment of bacteria to maintain or enhance their growth, although fully heterotrophic conditions will not allow their survival (Calderini et al., 2022). The abundance of *Cryptomonas* could be also related to nitrogen metabolism. In an experiment performed by Krustok et al. (2016) in municipal wastewater treating photobioreactors, this flagellate was able to grow to a higher concentration with nitrogen existing mostly as NH_4_-N.

Hence, contrary to the prokaryotic community, the decrease in edge density reveals that the eukaryotic community turned simpler and more specialized during the granular stage (Xiao et al., 2023). This simplification was also observed in the fewer co-occurrences between the Euk-2 and Prok-2 sub-communities in the bipartite network (79) compared with the Euk-1 and Prok-2 (178). The specialization of the eukaryotic community is reflected by the increasing abundance of *Rotifera* and *Sessilida,* groups adapted to granular sludge which provide the space where they can attach avoiding the washing out of the system (Li et al., 2013; Weber et al., 2007). Both, being filter feeders create water currents and ingest suspended prey and fine sludge particles more efficiently removing non-flocculated bacteria (Böhme et al., 2009; Li et al., 2013). Rotifers, more complex than protozoa, can also predate on these microorganisms limiting their populations (Li et al., 2013).

### High wash-out dynamics is not a requisite for granulation

Despite the long settling times used in the reactor, and thus applying a low wash-out regime to the biomass, granules started to emerge after 16 days and at day 112 they were fully developed. Washing out the non-granulated biomass is considered an important selection force for sludge granulation. But according to the results presented here, high wash-out rates is not a prerequisite for granulation to occur, although the process is accelerated considerably. By way of comparison, in a previous experiment using the same reactor set-up, but with a settling time of 2 minutes, granulated biomass dominated the reactor already after 25 days (Liébana et al., 2019). Granulation at low wash-out dynamics has also been reported by other researchers (Barr et al., 2010a; Dulekgurgen et al., 2003; Weissbrodt et al., 2013), even with a total retention of biomass in the reactor (Chen et al., 2017). However, when long settling times are applied, higher shear forces have been found necessary to achieve granulation (Chen and Lee, 2015; Zhou et al., 2014). These results suggest that other factors than short settling time may be more important for granulation, such as high hydrodynamic shear forces and feast-famine regimes. This opens the door to explore alternative strategies for granulation in different conditions, such as continuous operation (Liébana et al., 2018).

## Conclusions

Our findings provide insights in the successional patterns of micro-eukaryotes during granule formation and the interkingdom interactions of this population with the prokaryotic community, which has been neglected in the literature. Here, deterministic forces were important during sludge granulation, presumably caused by the acclimation of the microbial community to new environmental factors. Changes in the prokaryotic community structure, determined the successional patterns of the micro-eukaryotic communities. Although inter-kingdom interactions were shown to affect community succession during the whole experiment, during granule development random factors like the availability of settlement sites or drift acquired increasing importance.

