## Supplemental Material for "Community successional patterns and inter-kingdom interactions during granular biofilm development"

#Corresponding authors:

### Supplementary material

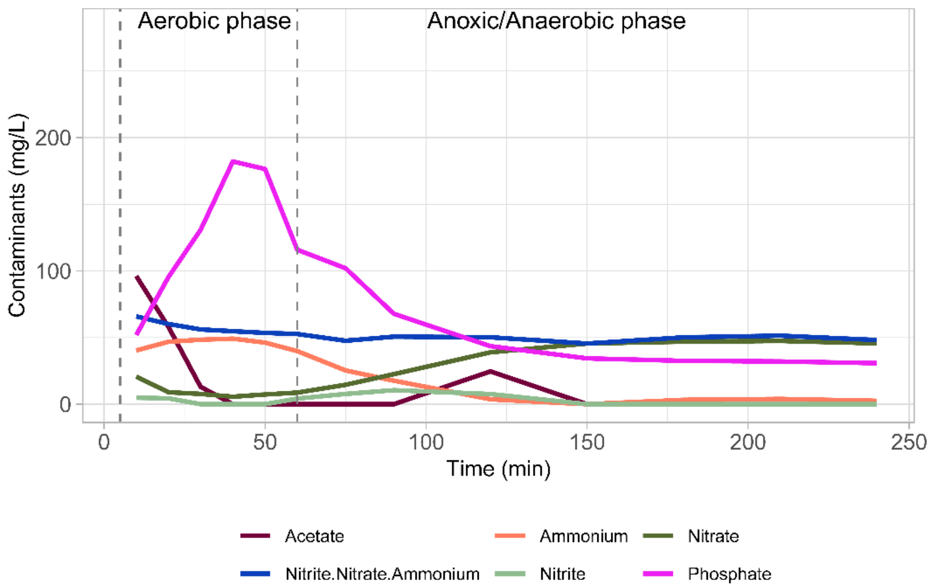


Figure S1. Cycle study results for acetate-C, nitrate-N, nitrite-N, ammonium-N, phosphate-P and total nitrogen expressed as the sum of nitrate, nitrite and ammonium.

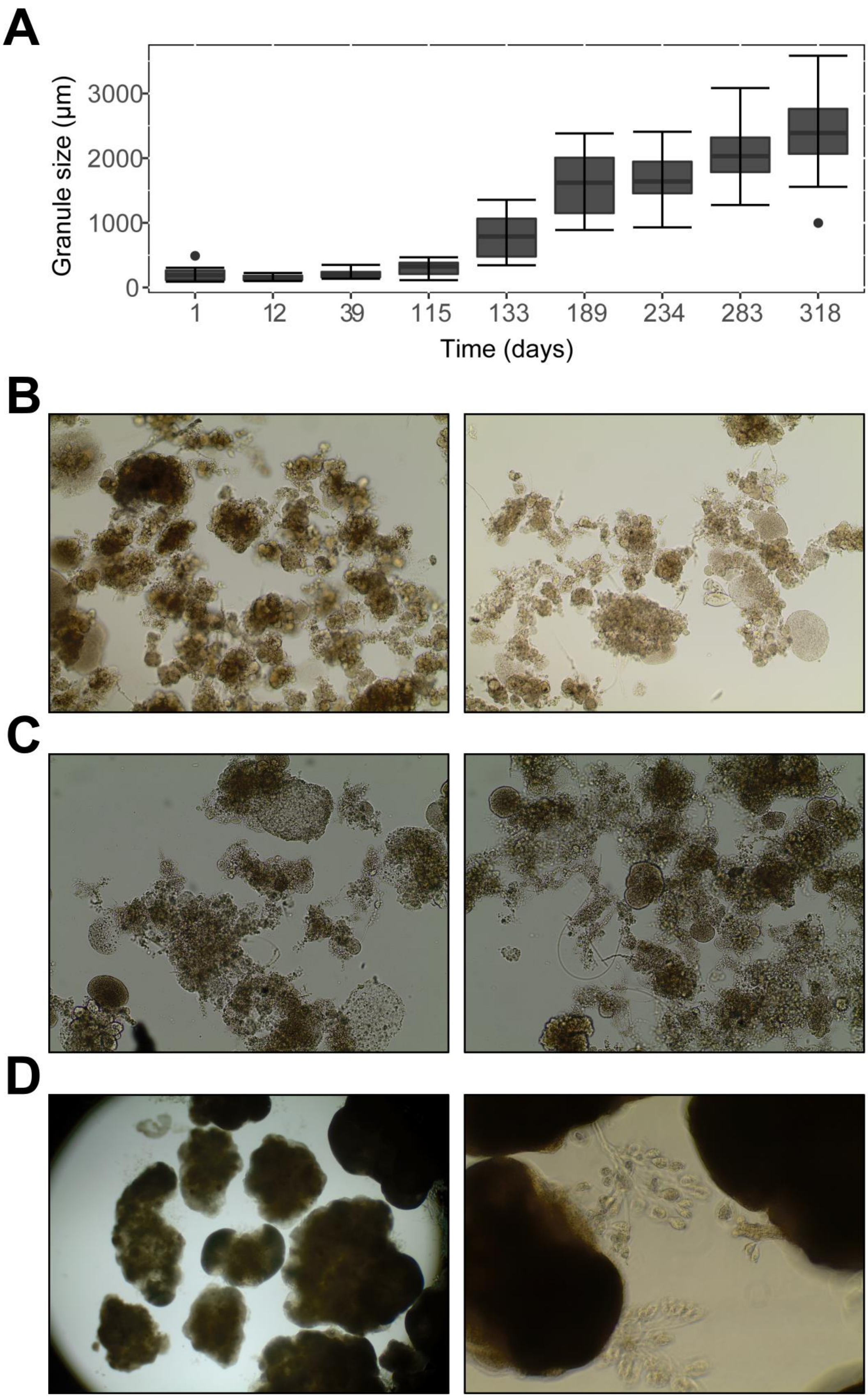


Figure S2. Particle size of the reactor sludge during experiment. A, granule size (µm). Microscopic image of sludge during the B, floccular; C, intermediate; and D, granular stage.

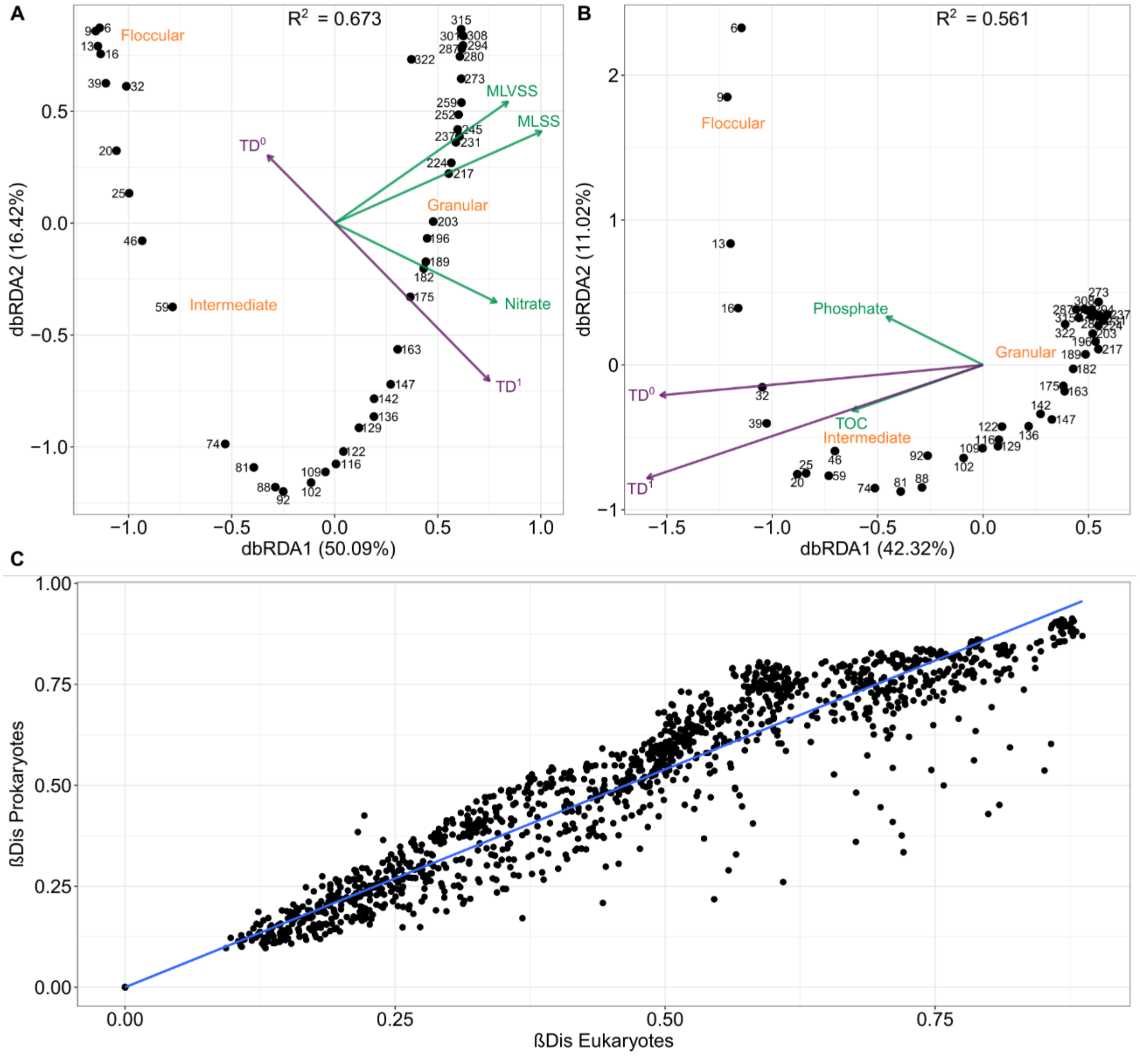


Figure S3. Constrained principal coordinates analysis (dbRDA) based on Bray-Curtis dissimilarity of square root transformed count tables. A, Prokaryotes. B, Eukaryotes. Significant variables ($p < 0.05$) were selected with an automated stepwise model (ordistep). Explained variance of each axis are indicated in parenthesis. The numbers refer to days of reactor operation. C, Correlation between prokaryotic and eukaryotic β-diversity, as measured by Bray-Curtis dissimilarity.

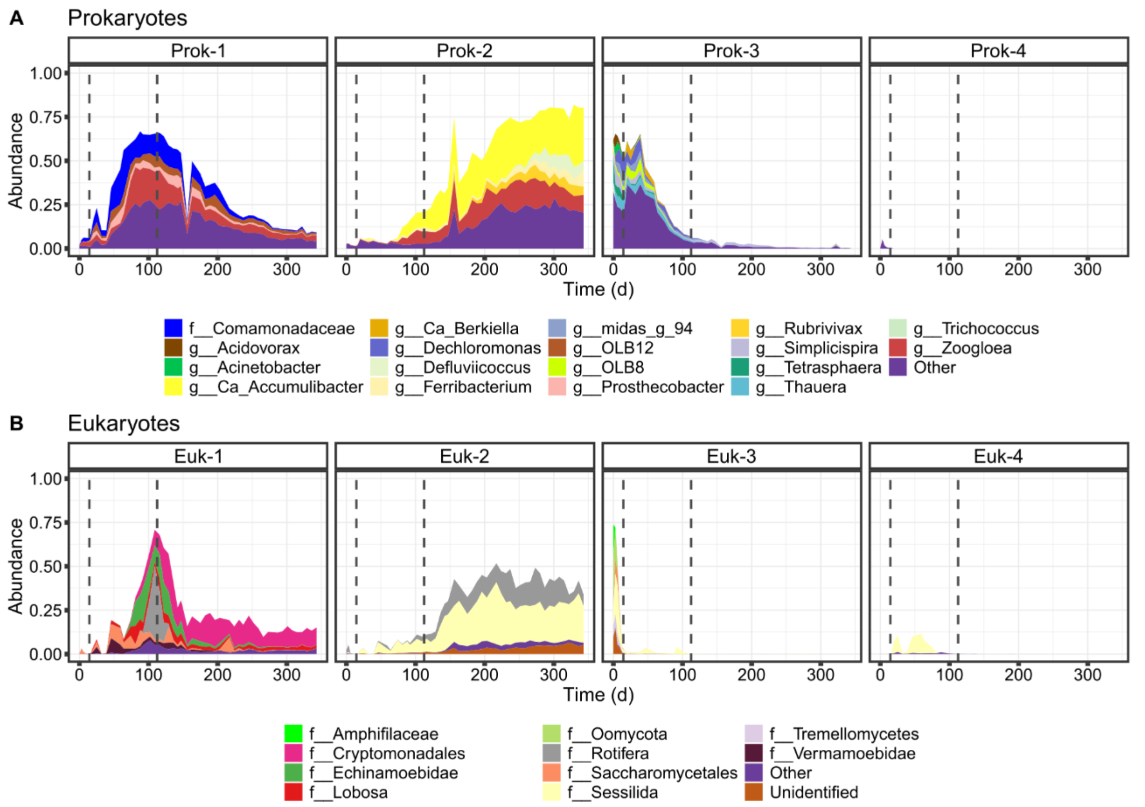


Figure S4. Taxonomic profiling of the modules detected in the microbial communities. A, taxonomic composition at the genus level of the modules detected in the prokaryotic co-occurrence metaweb. B, taxonomic composition at the family level of the modules detected in the eukaryotic co-occurrence metaweb. “Other”, includes genera with less than 5% abundance in at least one sample point; and “Unidentified”, taxonomically unassigned taxa.

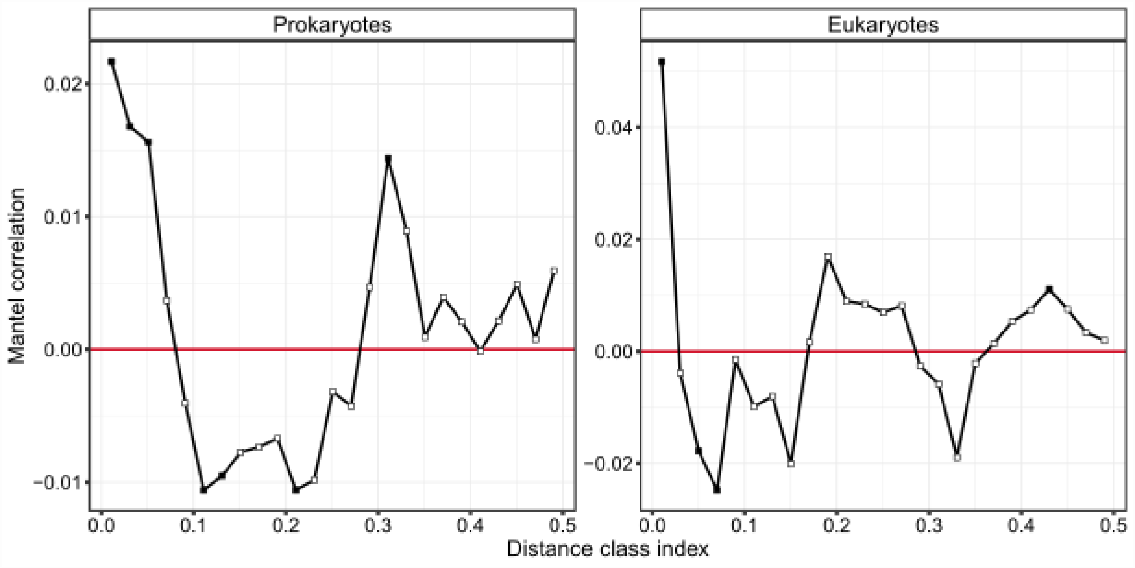


Figure S5. Mantel correlograms showing phylogenetic signal. Pearson correlation resulting from Mantel correlogram (999 permutations) between the ASV environmental (accounting for operational conditions and granulation stage) and phylogenetic distances for A, prokaryotic and B, eukaryotic communities. Significant correlations (solid squares) at low distances indicate phylogenetic signal in species ecological niches.

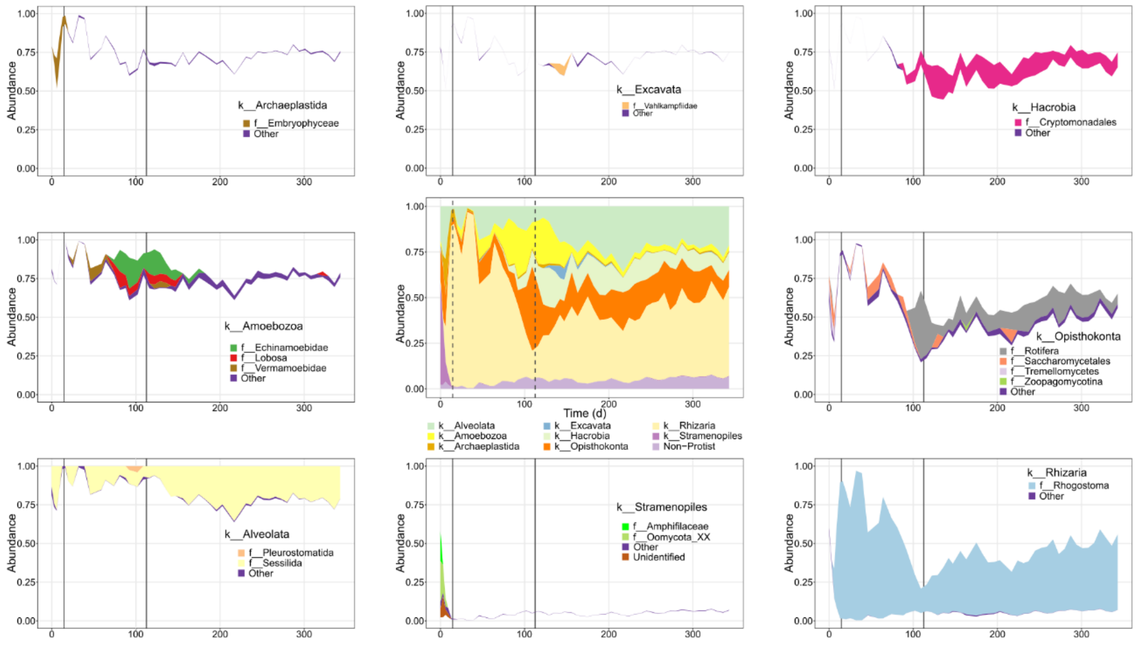


Figure S6. Temporal dynamics of eukaryotic taxonomic composition. Taxonomic distribution of the eukaryotic community at the superphylum and family levels. The centre panel represent the taxonomic profiling at the superphylum level and each panel next to it represent the taxonomic profiling at the family level of each superphylum.

Supplementary Table S1: Spearman's rank correlations between prokaryotic and eukaryotic community succession, taxonomic β-diversity between successive sample points, at diversity order of q.

| $\beta DisTD^0$ | $\beta DisTD^1$ | $\beta DisTD^2$ |
|---|---|---|
| 0.595 *** | 0.668 *** | 0.394 ** |

Asterisks denote the significance levels (***p-value < 0.001, **p-value < 0.01 and *p-value < 0.05).

Supplementary Table S2: Taxonomic and abundance information of the nodes from the core bipartite network.
